# Distracting Linguistic Information Impairs Neural Tracking of Attended Speech

**DOI:** 10.1101/364042

**Authors:** Bohan Dai, James M. McQueen, René Terporten, Peter Hagoort, Anne Kösem

## Abstract

Listening to speech is difficult in noisy environments, and is even harder when the interfering noise consists of intelligible speech as compared to unintelligible sounds. This suggests that the competing linguistic information interferes with the neural processing of target speech. Interference could either arise from a degradation of the neural representation of the target speech, or from increased representation of distracting speech that enters in competition with the target speech. We tested these alternative hypotheses using magnetoencephalography (MEG) while participants listened to a target clear speech in the presence of distracting noise-vocoded speech. Crucially, the distractors were initially unintelligible but became more intelligible after a short training session. Results showed that the comprehension of the target speech was poorer after training than before training. The neural tracking of target speech in the delta range (1–4 Hz) reduced in strength in the presence of a more intelligible distractor. In contrast, the neural tracking of distracting signals was not significantly modulated by intelligibility. These results suggest that the presence of distracting speech signals degrades the linguistic representation of target speech carried by delta oscillations.

## Introduction

Speech communication in everyday life often takes place in the presence of multiple talkers or background noise. In such auditory scenes, comprehension of target speech is often degraded due to interference from concurrent sounds (Bee and Micheyl, 2008; Cherry, 1953). Behavioral evidence specifically highlights that distracting sounds do not only mask the acoustics of the target speech input signal, but also interfere with its cognitive processing (Brungart et al., 2001; Woods and McDermott, 2015). Given that speech is a complex auditory signal that carries linguistic information, the interference between distracting sounds and target speech can occur at multiple levels of the speech processing hierarchy: interference could occur during the auditory analysis of speech, or at a later stage during the decoding of linguistic information (Evans and Davis, 2015; Hoen et al., 2007; Mattys et al., 2012; Rhebergen et al., 2005). In line with this prediction, interfering signals with different types of acoustic and/or linguistic information differently influence the perception and recall of target speech (Ellermeier et al., 2015; Wöstmann and Obleser, 2016). The comprehension of target speech is known to depend on the acoustic characteristics of the distracting sounds, for example, interference is stronger when target and distractor speech signals are spoken by talkers of the same sex than of different sexes (Brungart, 2001; Scott et al., 2004). On top of acoustic effects, the intelligibility of distracting speech alone is also a source of interference, such that the same noise-vocoded speech background impairs more strongly the comprehension of target speech when it is intelligible as compared to when it is unintelligible (Dai et al., 2017).

Distracting speech therefore interferes with the processing of abstract linguistic features of target speech. The neural origins of this interference, however, are still unclear. Interference could either arise from a degradation of the neural representation of the target speech, or from increased representation of distracting speech that enters in competition with the target speech. To test these alternative hypotheses, we observed the alignment between brain activity and speech dynamics, often called ‘neural tracking’ (Obleser and Kayser, 2019). Degree of neural tracking indicates how well each speech stream is segregated from the listening background (Ding and Simon, 2012a; Lakatos et al., 2008; Zion Golumbic et al., 2013). When listening to clear speech, neural oscillatory activity in the delta (1–4 Hz) and theta (4–8 Hz) ranges entrains to the dynamics of speech (Ahissar et al., 2001; Ding et al., 2016; Ding and Simon, 2012a, 2012b; Gross et al., 2013; Luo and Poeppel, 2012, 2007). In a noisy or multi-talker scene, both theta and delta neural oscillations primarily entrain to the dynamics of the attended speech (Ding and Simon, 2012a, 2012b; Mesgarani and Chang, 2012; Zion Golumbic et al., 2013). Yet, it is still under debate which aspects of speech are encoded in speech-brain tracking activity. Broadly speaking, this could be either acoustic features or higher-level language information (Ding and Simon, 2014; Kösem and Wassenhove, 2017). Recent work suggests that the different neural oscillatory markers link to distinct aspects of speech perception: theta tracking underlies speech sound analysis (Di Liberto et al., 2015; Kösem et al., 2016; Millman et al., 2014) while delta oscillations reflect higher-level linguistic processing, such as semantic and syntactic processing (Di Liberto et al., 2018; Ding et al., 2016).

Here, we examined whether competing linguistic information influences the neural tracking of target and ignored speech in a cocktail-party setting. In order to isolate the linguistic effect from the effect of acoustic competition between the target and distracting speech, we used a A-B-A training paradigm in which the intelligibility of the distracting stimulus was manipulated while its acoustic properties were kept constant (Dai et al., 2017). Participants performed a dichotic listening task twice, in which they were asked to repeat a clear speech signal while noise-vocoded (NV) speech was presented as distractor (Fig. 1A). In between the two sessions, participants were trained to understand the interfering NV distractor (Fig. 1B). We compared behavioral performance (accuracy in the repetition of the target speech) and MEG oscillatory activity between the two dichotic listening sessions. Our main prediction was that distracting speech would impair more strongly target speech comprehension and neural tracking when it gained intelligibility. Based on previous reports (Ahissar et al., 2001; Ding and Simon, 2013; Doelling et al., 2014; Peelle et al., 2012), we also predicted that neural tracking of distracting speech would be stronger when it is more intelligible.

**Figure 1.**
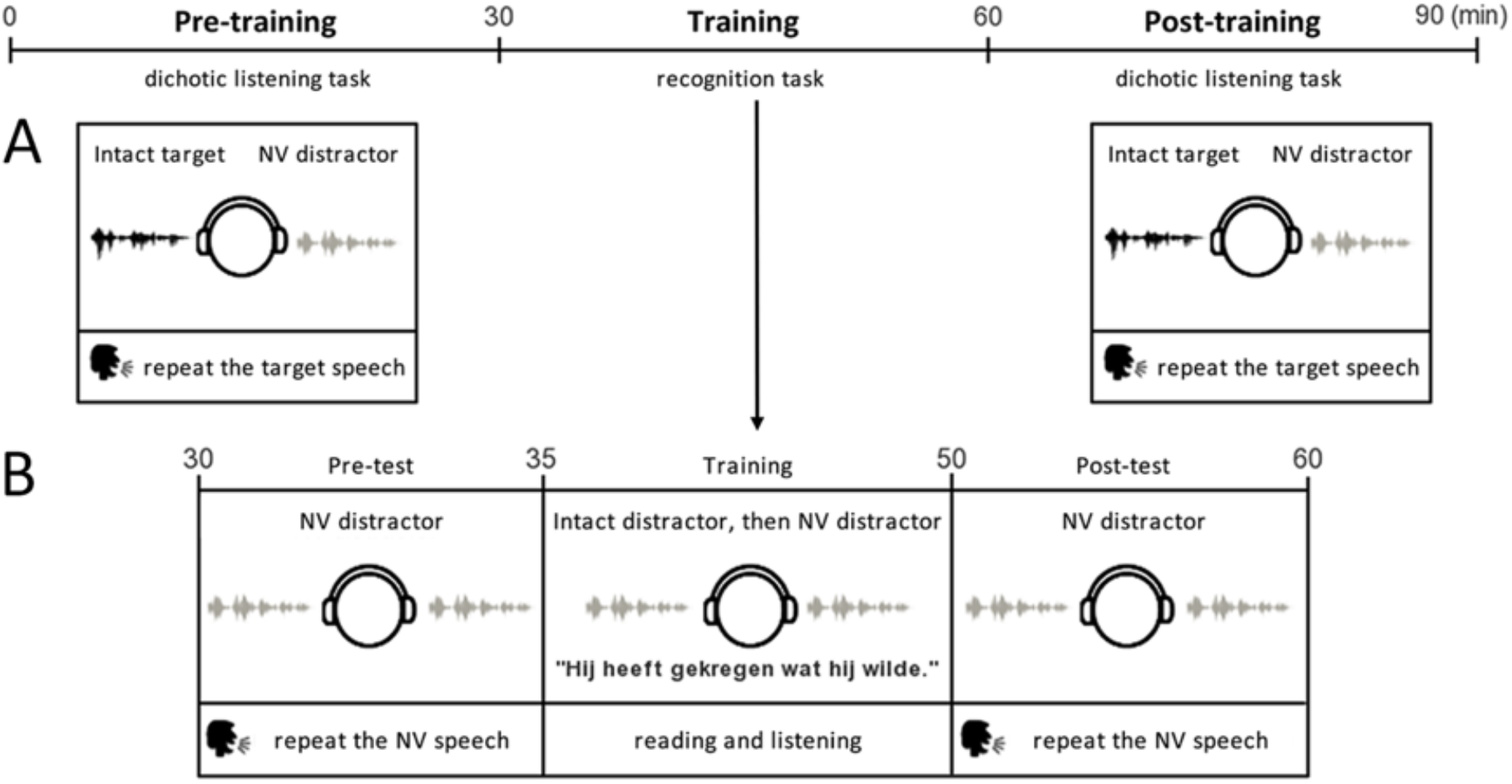
Experimental design. The experiment consisted of three phases. In the first and third phases of the experiment, participants performed a dichotic listening task (A), in between the two dichotic listening tasks participants were trained to understand 4-band NV speech (B). (A) During the dichotic listening tasks, participants listened to the presentation of one intact target with one NV distractor (either 2- or 4-band NV speech) and were asked to repeat the intact target speech. (B) During the training of the 4-band NV sentences, participants listened to the distractor once in the intact and then once in the NV version. At the same time, they read the text of the sentences on the screen. We tested the intelligibility of the 4-band NV sentences before (pre-test) and after (post-test) the training by asking participants to listen to and repeat the NV sentences. Participants were also familiarized with 2-band NV sentences in pre-test and post-test to assess their intelligibility.

## Materials and Methods

### Participants

Twenty-seven participants (13 women, 18~33 years old, mean age: 23.5 ± 3.9 years) took part in the study. All were right-handed native Dutch speakers with normal hearing (self-reported through the questionnaire in the participant recruit system of Donders Institute for Brain, Cognition and Behaviour, Radboud University). Two participants were rejected, one due to malfunctioning of the MEG system and one because the participant did not finish the task. The analyses are thus based on the data of 25 participants (12 women, 18~33 years old, mean age: 23.5 ± 4.0 years). All participants gave their informed consent in accordance with the Declaration of Helsinki, and the local ethics committee (CMO region Arnhem-Nijmegen).

### Stimuli

As in our previous study (Dai et al., 2017), the stimuli were selected from a corpus with meaningful conversational Dutch sentences (e.g., ‘Mijn handen en voeten zijn ijskoud’, in English: ‘My hands and feet are freezing’), digitized at a 44,100 Hz sampling rate and recorded at the VU University Amsterdam (Versfeld et al., 2000) by a male or female speaker. Each sentence consisted of 5-8 words. Each speech stimulus consisted of a combination of two semantically independent sentences of the corpus uttered by the same speaker, separated by a 300-ms silence gap (average duration = 4.15 ±0.13 s).

The target speech stimuli consisted of 384 intact sentence pairs spoken by one of the two speakers (half of the trials were from the male speaker and half were from the female speaker). The distracting speech stimuli were NV versions of 48 different sentence pairs taken from the same corpus and spoken by the other speaker (i.e., a speaker of opposite sex). Noise-vocoding (Davis et al., 2005; Shannon et al., 1995) was performed with Praat software (Version: 6.0.39 from http://www.praat.org), using either 4 (main condition) or 2 (control) frequency bands logarithmically spaced between 50 and 8000 Hz. In essence, the noise-vocoding technique parametrically degrades the spectral content of the acoustic signal (i.e., the fine structure) but keep the temporal information largely intact.

### Procedure

The main experiment was similar to our previous study (Dai et al., 2017). The experimental design was implemented using Presentation software (Version 16.2, www.neurobs.com). The experiment included three phases: pre-training, training, and post-training. In the pre- and post-training phases, the participants performed the dichotic listening task. Each trial consisted of the presentation of the target speech with the interfering NV speech. The two signals were delivered dichotically to the two ears. We used here a dichotic listening task to intentionally alleviate the effects of energetic masking. The target side (left or right) for a particular trial was pseudo-randomly defined: half of the trials had the target on the left ear and half had it on the right ear. The stimuli were presented at a comfortable listening level (70 dB) by MEG-compatible air tubes in a magnetically shielded room. The signal-to-noise ratio (SNR) was fixed on −3 dB based on the results of our previous study, −3dB SNR being the condition in which we observed the strongest masking effects of intelligible distractor signals. The participants were requested to listen to the presentation of one intact speech channel and one unintelligible NV speech channel and to pay attention to the clear speech only, which was hence the target speech. This instruction was well understood by the participants as the NV speech was strongly distorted and sounded very different from clear speech. The participants were not told which ear the target speech would be presented to. After the presentation, the participant’s task was to repeat the sentences of the target speech. Participants’ responses were recorded by a digital microphone with a sampling rate of 44,100 Hz. The distracting speech consisted of 24 sentences of 4-band NV speech and 24 sentences of 2-band NV speech. Each dichotic listening task comprised 192 trials total. The target speech differed across trials and all conditions (hence a target stimulus was only presented once during the whole experiment), while NV distracting stimuli were repeated four times, for a total of 96 trials per NV condition. The 24 same NV sentences presented during training were used during pre- and posttraining sessions. The ear of presentation of the target signals, and the type of NV signals (4-band, 2-band), were randomized across trials. Each dichotic listening task was 30 min long.

In the training phase, participants were trained to understand the 4-band NV speech. The training phase included three parts: (a) pre-test: the participants were tested on their ability to understand the 24 4-band NV stimuli and 24 2-band NV stimuli used in the dichotic listening task as distracting signals; they were presented with the interfering speech binaurally and were asked to repeat it afterwards; (b) training on 4-band NV speech: they were presented one token of an intact version of an NV stimulus followed by one token of the NV version of that stimulus; at the same time, they could read the content of the NV speech on the screen; 2-band NV speech was not trained; (c) post-test: they performed the intelligibility test again. Crucially, the 4-band NV speech were initially poorly intelligible but could be better understood after training (Dai et al., 2017; Davis et al., 2005; Sohoglu and Davis, 2016). Hence, during the pre- and post-training phases, the NV speech would have the same acoustic information but would have different levels of intelligibility. In total, the training phase took 30 min.

### Behavioral analysis

The intelligibility of speech was measured by calculating the percentage of correct content words (excluding function words) in participants’ reports for each trial. Words were regarded as correct if there was a perfect match (correct word without any tense errors, singular/plural form changes, or changes in sentential position). The percentage of correct content words was chosen as a more accurate measure of intelligibility based on acoustic cues than percentage correct of all words, considering that function words can be guessed based on the content words (Brouwer et al., 2012). For the training phase, we performed a two-way repeated-measures ANOVA with noise vocoding (trained 4-band and untrained 2-band) and time (pre- and post-training) as factors. For the dichotic listening tasks, a three-way repeated-measures ANOVA was performed to assess the contribution of three factors: noise vocoding (4-band and 2-band), time (pre-training and post-training), and side (left target/right distractor and right target/left distractor). In our post hoc sample t-tests, we compensated for multiple comparisons with Bonferroni correction.

### MEG Data Acquisition

MEG data were recorded with a 275-channel whole-head system (CTF Systems Inc., Port Coquitlam, Canada) at a sampling rate of 1200 Hz in a magnetically shielded room. Data of two channels (MLC11 and MRF66) were not recorded due to channel malfunctioning. Participants were seated in an upright position. Head location was measured with two coils in the ears (fixed to anatomical landmarks) and one on the nasion. To reduce head motion, a neck brace was used to stabilize the head. Head motion was monitored online throughout the experiment with a real-time head localizer and if necessary corrected between the experimental blocks.

### MEG Data preprocessing

Data were analyzed with the FieldTrip toolbox (fieldtrip-20171231) implemented in MATLAB (Oostenveld et al., 2011). Trials were defined as data between 500 ms before the onset of sound signal and 4, 000 ms thereafter. Three steps were taken to remove artifacts. Firstly, trials were rejected if the range and variance of the MEG signal differed by at least an order of magnitude from the other trials of the same participant. On average, 14.1 trials (3.67%, SD = 7.8) per participant were rejected via visual inspection. Secondly, data were down-sampled to 100 Hz and independent component analysis (ICA) was performed. Based on visual inspection of the ICA components’ time courses and scalp topographies, components showing clear signature of eye blinks, eye movement, heartbeat and noise were identified and removed from the data. On average, per participant 9.8 components (SD = 2.6) were rejected (but no complete trials). Finally, 9.6 trials (2.5%, SD = 4.4) with other artifacts were removed via visual inspection like the first step, resulting in an average of 360.3 trials per participant (93.83%, each condition: ~90 trials). Subsequently, the cleaned data were used for further analyses.

### MEG analysis

A data-driven approach was performed to identify the reactive channels for sound processing. The M100 (within the time window between 80ms and 120ms after the first word were presented) response was measured on the data over all experimental conditions, after planar gradient transformation (Oever et al., 2017).We selected the 18 channels with the relatively strongest response at the group level on each hemisphere, and the averages of these channels were used for all subsequent analysis. The locations of the identified channels cover the classic auditory areas.

Speech-brain coherence analysis was performed on the data within the region of interest after planar gradient projection. Spectral analysis was performed using ‘dpss’ multi-tapers with a ± 1 Hz smoothing window of the speech envelopes, and of the neural times series epoched from 500 epochs were removed to exclude the evoked response to the onset of the sentence. To match trials in duration, shorter trials were zero-padded up to 3.4s (the max length of the signal) for both target speech and distracting speech. The speech-brain coherence was measured at different frequencies (1 to 15 Hz, 1 Hz step). Finally, the coherence data were projected into planar gradient representations. Then data were averaged across all trials and the strongest 36 channels defined by our auditory response localizer. Following the same method, we calculated the surrogate of speechbrain coherence (as control condition) by randomly selecting the neural activity of one trial and the temporal envelope of speech of another trial. For the investigation of our main hypotheses, we restricted the speech-brain coherence analyses to delta band (1–4 Hz) and theta band (4–8 Hz) activity; these frequency bands were chosen based on the previous literature (Ding and Simon, 2012b, 2012a; Gross et al., 2013; Kösem and Wassenhove, 2017; Luo and Poeppel, 2012). Restricting our analysis to frequency bands of interests and regions of interest (auditory areas) allowed us to perform a three-way repeated-measure ANOVA with frequency (delta, theta), speech type (target, distractor), and data (data, surrogate) as factors.

We repeated the same analysis described above to quantify the speech-brain coherence for each condition, and the averaged speech-brain coherences of the strongest 18 channels on each hemisphere in the delta and theta bands were calculated. We tested the target and distractor speechbrain coherence in the delta and theta range using a four-way repeated measure ANOVA with factors noise vocoding (4-band, 2-band), time (pre-training, post-training), side (left target, right target), and hemisphere (left, right). The relative coherence change of training was calculated based on the following formula: relative change = (Coh_post_ – Coh_pre_) / (Coh_post_ + Coh_pre_). Afterward, the correlation of coherence change and behavioral change was calculated based on Spearman’s rho.

### MRI data acquisition and source reconstruction analysis

Anatomical MRI scans were obtained after the MEG session using either a 1.5 T Siemens Magnetom Avanto system or a 3 T Siemens Skyra system for each participant (except for one participant whose data was excluded from source reconstruction analysis). The co-registration of MEG data with the individual anatomical MRI was performed via the realignment of the fiducial points (nasion, left and right pre-auricular points). Lead fields were constructed using a single shell head model based on the individual anatomical MRI. Each brain volume was divided into grid points of 1 cm voxel resolution, and warped to a template MNI brain. For each grid point the lead field matrix was calculated. The sources of the observed delta and theta speech-brain coherence were computed using beamforming analysis with the dynamic imaging of coherent sources (DICS) technique to the coherence data.

## Results

### Intelligible NV speech interfered more with target speech’s understanding

Two types of NV speech were used as distractors in the dichotic listening task: either 4-band or 2-band NV speech segments. In the training phase, participants were trained to understand 4-band NV speech, while 2-band NV speech was presented but not trained. Hence, 2-band NV speech served as control distractors that would not improve in intelligibility with training. To make sure the training was efficient in improving the intelligibility of 4-band NV speech, we compared the participants’ comprehension of the NV signals before and after training. Consistent with previous findings (Dai et al., 2017), and as shown in Fig. 2A, the training significantly improved the perception of 4-band NV speech. A two-way repeated-measure ANOVA showed that the main effects of noise vocoding (2-band *vs.* 4-band) and time (pre- *vs.* post-training) were significant (noise vocoding: (*F*(1, 24) = 217.78, *p* < 0.001, *η2* = .90; time: *F*(1, 24) = 219.07, *p* < 0.001, *η2* = .90). Crucially, a significant interaction between noise vocoding and time was observed (*F*(1,24) = 262.94, *p* < 0.001, *η2* = .92), meaning that the intelligibility of 4-band NV speech was significantly improved compared to that of 2-band NV speech (4-band(post-pre) vs. 2-band(post-pre): *t* (24) = 16.22, *p* < 0.001, Cohen’s *d* = 3.20). After training, 4-band NV sentences had a score of 52.69 ± 3.35% recognition accuracy (31.70 ± 2.03% improvement during training; values here and below indicate mean ± SEM), while 2-band NV sentences remained mostly unintelligible with a score of 1.97 ± 0.52% recognition accuracy (1.06 ± 0.35% improvement during training).

**Figure 2.**
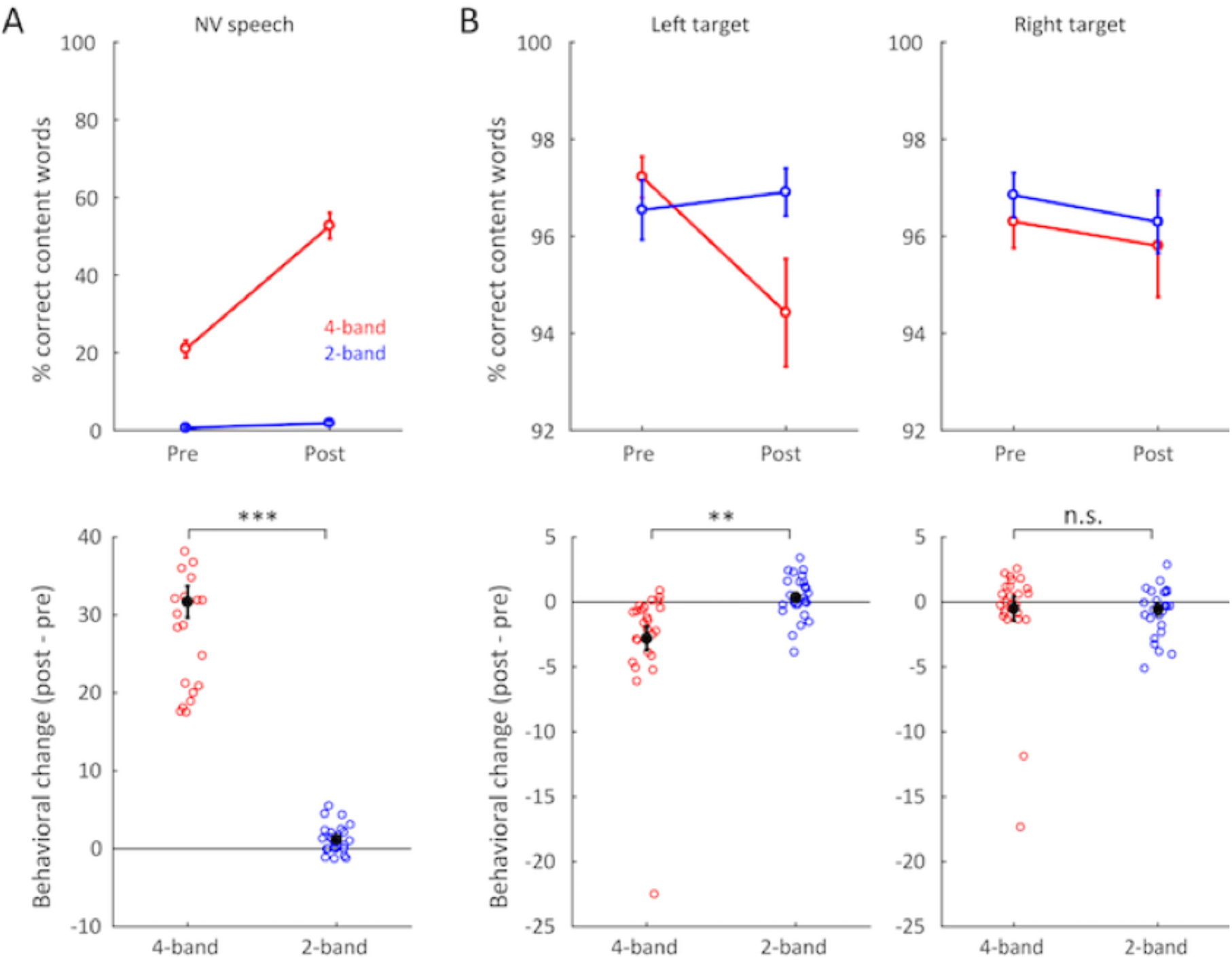
Behavioral results. (A) Intelligibility of NV speech during the training phase. The intelligibility of trained 4-band (red) significantly improved by more than 30% with training, while untrained 2-band (blue) NV speech remained mostly unintelligible post-training. Top panel indicates the grant average of performance in two training phase. Bottom panel indicates the individual change through training. Each circled dot corresponds to one participant. The filled black dot represents the average. Error bars indicated standard error of the mean. (B) Intelligibility of target speech in the dichotic listening tasks. The intelligibility of target speech decreased after training when presented in competition with the 4-band NV speech (red), i.e. when distracting NV speech was more intelligible. The intelligibility of target speech was not significantly affected by training when presented in competition with the untrained 2-band NV speech (blue). This is only observed when the target speech is delivered to the left ear and the distractor to the right ear (left panel) and not when the target is delivered to the right ear and the distractor to the left ear (right panel). Top panel indicates the grant average of performance in two training phase. Bottom panel indicates the individual change through training. Each circled dot corresponds to one participant. The filled black dot represents the average. Error bars indicated standard error of the mean. *** *p* < 0.001, ** *p* < 0.01, and n.s., not significant.

The training thus efficiently improved the intelligibility of 4-band NV speech. We then investigated if the change in the intelligibility of the distractor interfered with the comprehension of the target speech during dichotic listening. To assess the magnitude of increased interference, we measured the accuracy of target speech recognition in the two dichotic listening tasks (Fig. 2B). A three-way repeated-measure ANOVA was performed (time (pre-training, post-training), noise vocoding (trained 4-band, untrained 2-band), and side of target presentation (left target, right target)). We observed a significant main effect of noise vocoding (*F*(1, 24) = 6.44, *p* < 0.05, *η2* = .21), and a significant interaction of the three factors (*F*(1, 24) = 8.22, *p* < 0.01, *η2* = .26), which revealed that the change of interference depended on which ear the target speech was delivered. A closer look at the data showed that target speech comprehension decreased after training when the target speech was presented to the left ear (i.e., when the distractor NV signal was presented to the right ear). There was a significant interaction between noise vocoding and time (*F*(1, 24) = 8.12, *p* < 0.01, *η2* = .28): the 4-band NV speech interfered more strongly with target speech comprehension after training than before training (pre = 97.22 ± 0.42%; pot = 94.42 ± 1.11%; post vs pre: *t* (24) = −3.08, *p* < 0.01), while 2-band NV speech had a similar masking effect before and after training (pre = 96.54± 0.61%; post = 96.91 ± 0.49%; post vs pre: *t* (24) = −0.93, *p* = 0.362). As previously shown (Dai et al., 2017), these findings suggest that the increased intelligibility of 4-band NV speech acquired during training generates more interference in the processing of the target speech and decreases its comprehension. When the target speech was delivered to the right ear (and the distractor to the left ear), no effect of training was observed (interaction of noise-vocoding and time: *F*(1, 24) = 0.003, *p* = 0.954, *η2* = .0001; 4-band: pre = 96.30 ± 0.54%, post = 95.80 ± 1.05; 2-band: pre = 96.85 ± 0.46%, post = 96.29 ± 0.65%). This is in line with previous studies showing a right ear advantage in the recognition of dichotically presented speech. Effects of ear of presentation were not reported in Dai et al. (2017), but additional analysis of those data revealed a similar (but not statistically significant) asymmetric pattern.

### Neural tracking of target speech and distracting speech

In the first stage of MEG analysis, we inspected the speech-brain coherence for both target speech and distracting speech (Fig. 3). We focused on both delta (1–4 Hz) and theta (4–8 Hz) tracking of speech, as both frequency ranges are deemed relevant for speech processing (Ding et al., 2016; Ding and Simon, 2012b, 2012a; Giraud and Poeppel, 2012; Gross et al., 2013; Kösem and Wassenhove, 2017; Luo and Poeppel, 2012, 2007; Park et al., 2015). Coherence data were computed from the 36 channels (18 channels in each hemisphere) that produced the strongest auditory evoked M100 responses (Fig. 3), and were first averaged across all conditions (i.e., across the pre- and post-training sessions and across distractor type) and all sensors. A three-way repeated-measure ANOVA was performed (frequency (delta, theta), speech type (target, distractor), and data (data, surrogate)). Compared to surrogate coherence between neural oscillations and speech envelope, we observed stronger target- and distractor-brain coherence in both delta and theta ranges (main effect of data: (*F*(1, 24) = 56.22, *p* < 0.0001, *η2* = .70; Fig. 3A-B). A main effect of frequency was observed as well (*F*(1, 24) = 194.64, *p* < 0.0001, *η2* = .89), showing stronger speech-brain coherence in the delta range than in the theta range (Fig. 3A-B). Overall, neural tracking of target speech was stronger than that to distracting speech (main effect of speech type: (*F*(1, 24) = 54.98, *p* < 0.0001, *η2* = .69, interaction of data and speech type (*F*(1, 24) = 79.89, *p* < 0.0001, *η2* = .77, post-hoc tests: data target > data distractor, *p* < 0.0001; data target > surrogate target, p<0.0001; data distractor > surrogate distractor, p<0.0001; surrogate target vs. surrogate distractor, *p* = 1, Fig. 3A-B). As we used a dichotic listening task, speechbrain coherence was stronger on the contralateral side to the ear of presentation, for both target and distracting speech in both delta and theta ranges (Fig 3C-D; interaction side*hemi: delta tracking of target speech: *F*(1, 24) = 16.26, *p* < 0.001, *η2* = 0.40; delta tracking of distracting speech: *F*(1, 24) = 8.7, *p* =0.007, *η2* = .27; theta tracking of target speech: *F*(1, 24) = 36.46, *p* < 0.001, *η2* = .60; theta tracking of distracting speech: *F*(1, 24) = 22.64, *p* < 0.001, *η2* = .49). However, speech-brain coherence for both target and distracting speech was also observed on the ipsilateral side, suggesting that both signals were processed bilaterally even in a dichotic listening task.

**Figure 3.**
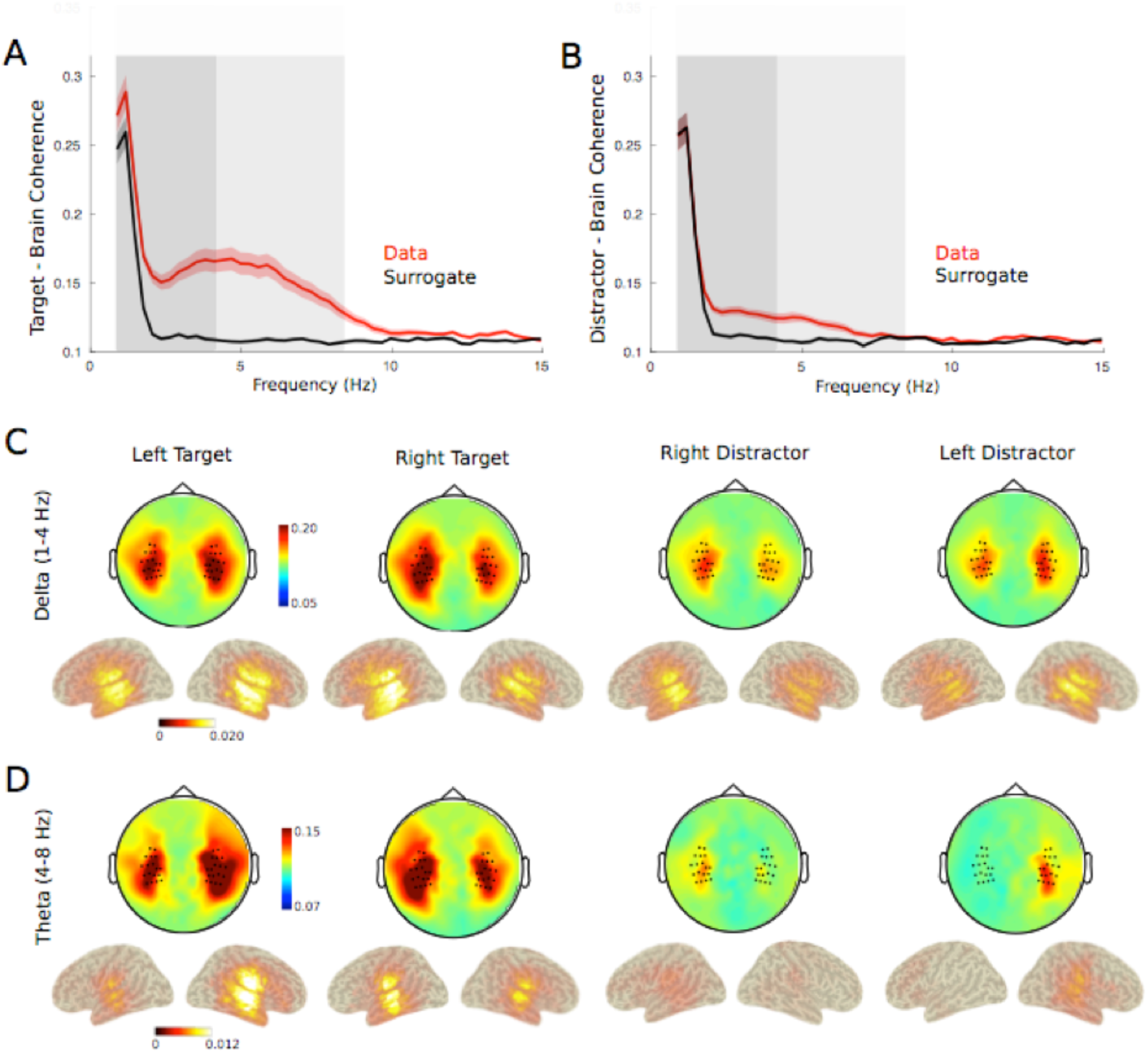
Neural tracking of target and distracting speech in both dichotic listening tasks. The top panel respectively show neural tracking of (A) target and (B) distracting speech from 1 to 15 Hz in selected temporal channels. The red line is the real speech-brain coherence data, while the black line is the control data which was calculated by randomly combining brain and speech data from different trials (see methods for details). The dark and light grey shadows mark respectively the delta and theta ranges as defined in our study. Topographies and source reconstruction of delta (C) and theta (D) tracking of target and distracting speech. Black dots in topographies mark the selected channels for the region-of-interest speech-brain coherence analyses. Speech-brain coherence is stronger in the auditory regions contralateral to the ear of presentation of the stimulus, though brain responses are observed bilaterally.

### Neural tracking of target speech was modulated by the intelligibility of the distracting speech

We then tested the effect of training on speech-brain coherence in the delta and theta ranges. Specifically, we expected the neural analysis of target speech (as reflected by speech-brain tracking) to be more impaired in the presence of an intelligible distractor. Hence, we predicted speech-brain coherence to target speech to become weaker after training, and this only for the 4-band NV distractor condition as the effect of training was limited to this type of distracting signal. This prediction was supported by target-brain coherence: delta tracking of target speech was reduced after training when distractor signals were 4-band NV speech, while the 2-band NV speech condition did not show this change (Fig. 4, interaction between noise-vocoding and time (*F*(1, 24) = 5.01, *p* = 0.035, *η2* = 0.17), post-hoc effects: 4-band, post vs. pre,*p* < 0.001; 2-band, post vs. pre: *p* = 0.60, Bonferroni corrected for multiple comparisons). As reported earlier, the four-way repeated-measure ANOVA showed a significant interaction effect between side and hemisphere (*F*(1, 24) = 16,26, *p* < 0.001, *η2* = 0.40), highlighting that target speech – brain coherence was stronger in the contralateral hemisphere. Yet, the effect of distractor’s intelligibility was not significantly affected by the side of target speech presentation, nor by the hemisphere of interest as no other significant interactions were found (interaction among hemisphere, NV and time: (*F*(1, 24) = 2.72, *p* = 0.11, *η2* = 0.10); interaction among side, NV and time: (*F*(1, 24) = 0.32, *p* = 0.58, *η2* = 0.01); interaction among hemisphere, side, NV and time: (*F*(1, 24) = 3.29, *p* = 0.08, *η2* = 0.12)). These results suggest that delta tracking of target speech in both hemispheres declined, regardless of the ear of presentation of the two speech signals.

**Figure 4.**
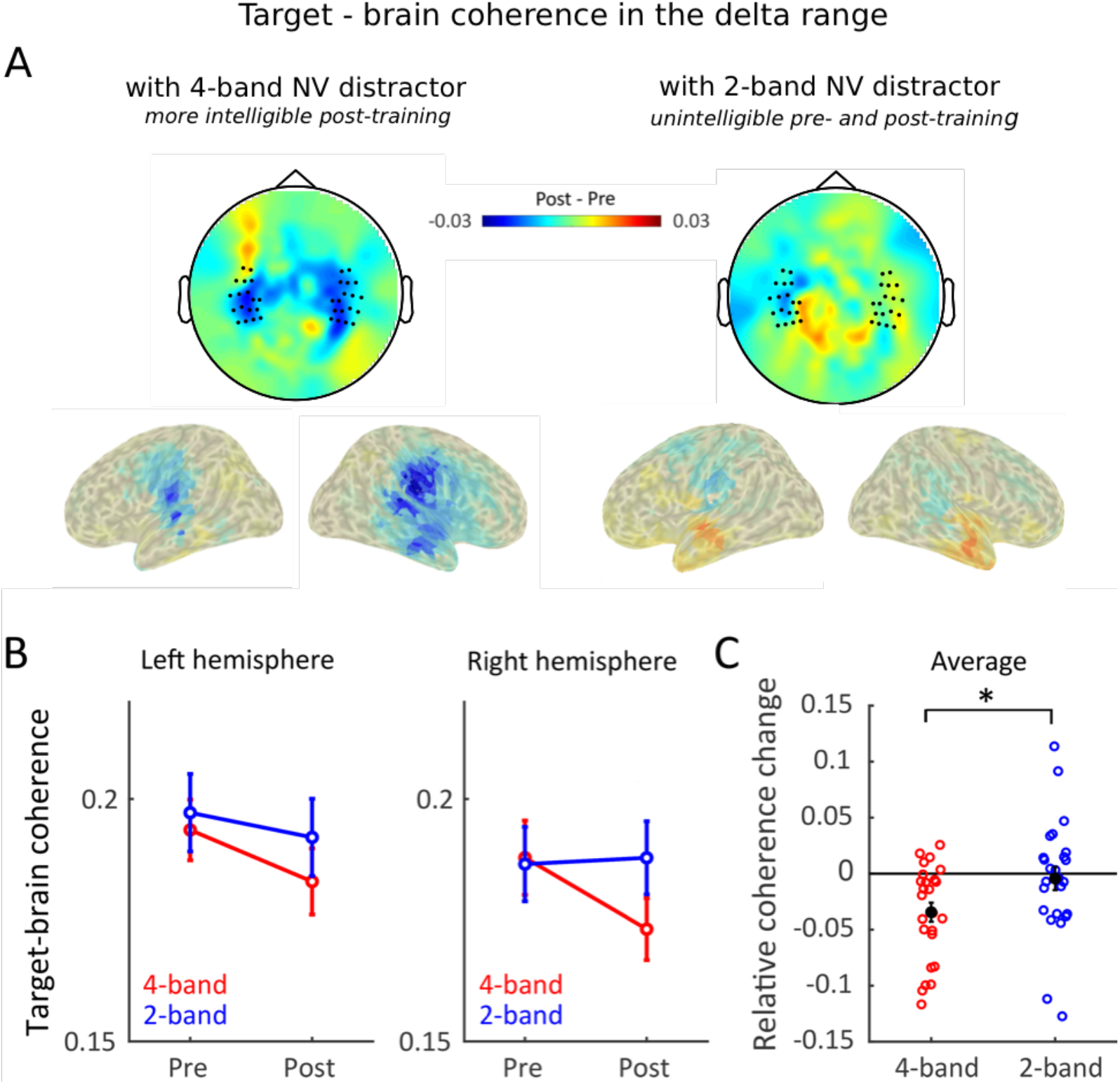
Delta tracking of target speech reduced when the distracting NV speech gained intelligibility. (A) Topographies and source reconstruction of delta tracking changes (Post-training minus Pre-training). A significant reduction in delta tracking of target was observed in bilateral temporal lobes after training when it was presented in competition with a distracting 4-band NV speech, i.e. when the distracting speech had gained intelligibility via training (left panel). No significant effects of training were observed when the target speech was presented with unintelligible 2-band NV distracters (right panel) (B) Delta tracking of target speech averaged across selected channels in each hemisphere, when in competition with distracting 4-band NV speech (red) or 2-band NV speech (blue). Error bars indicated standard error of the mean. (C) Relative Coherence change before and after training. Data is pooled across hemispheres as the effect of intelligibility was not significantly different across left and right temporal regions. Each circled dot corresponds to one participant. The filled black dot represents the average. Error bars indicated standard error of the mean. * *p* < 0.05.

To test whether the behavioral reduction was associated with the neural changes, we correlated the relative changes of target speech-brain tracking between pre- and post-training sessions (entrainment change = (Coh_post_ – Coh_pre_) / (Coh_post_ + Coh_pre_)) with the absolute behavioral change. A significant correlation between the relative change of delta tracking of target speech in the left hemisphere and the behavioral reduction of reporting target speech was observed, but only when target speech was delivered to the left ear *(rho* = 0.60, *p* = 0.008). However, this correlation was not significant after exclusion of one outlier participant (Fig. S1).

In contrast to delta oscillations, target-brain coherence in the theta range was not significantly affected by the intelligibility of the distractor (Fig. 5, interaction NV* Time: *F*(1, 24) = 0.43, *p = 0.52, η2* = 0.017). Theta tracking of target speech did show a significant interaction of the factors time, noise-vocoding and side of presentation (*F*(1, 24) = 6.05, *p = 0.02, η2* = 0.20). However, the post-hoc tests examining the interaction yielded no significant difference.

**Figure 5.**
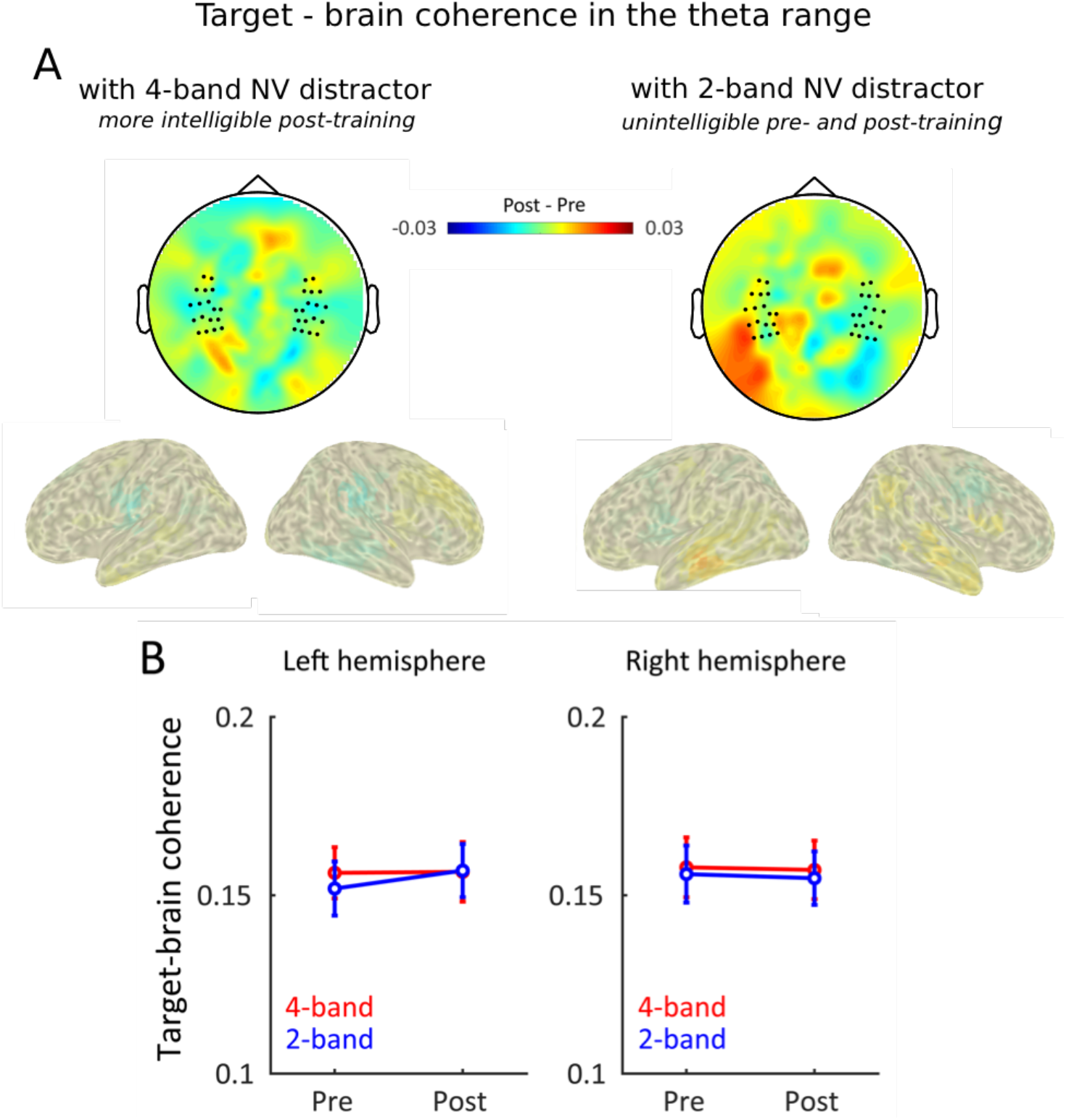
Theta tracking of target speech showed no significant change when the distracting NV speech gained intelligibility. (A) Topographies and source reconstruction of changes in speech-theta tracking (Post-training minus Pre-training). No significant changes of training were observed for both 4- and 2-band NV conditions. (B) Theta tracking averaged across selected channels in each hemisphere, when in competition with distracting 4-band NV speech (red) or 2-band NV speech (blue). Error bars indicated standard error of the mean.

### Neural tracking of distracting speech was not modulated by its intelligibility

We also tested whether the increased intelligibility of the distractor had an effect on the distractor-brain coherence. As previous studies suggested that speech-brain tracking is stronger for intelligible signals (Peelle et al. (2012), though see Millman et al. (2014) and Zoefel and VanRullen (2016)), we asked whether speech-brain coherence to distracting signals would increase after training for the intelligible 4-band NV distractor sentences. However, this effect was not present in the data. Distractor-brain coherence in the delta frequency was overall reduced after training compared to before training (Fig. 6, main effect of time: *F*(1, 24) = 38.42,*p* < 0.0001, *η2* = .62) irrespective of the type of distractor (2-band or 4-band NV speech, interaction NV*Time: *F*(1, 24) = 0.09, *p* = 0.76, *η2* = 0.00). The reduction of delta tracking of distracting speech can thus not be attributed to the training or the degree of intelligibility of the distractor, but may relate to habituation effects.

**Figure 6.**
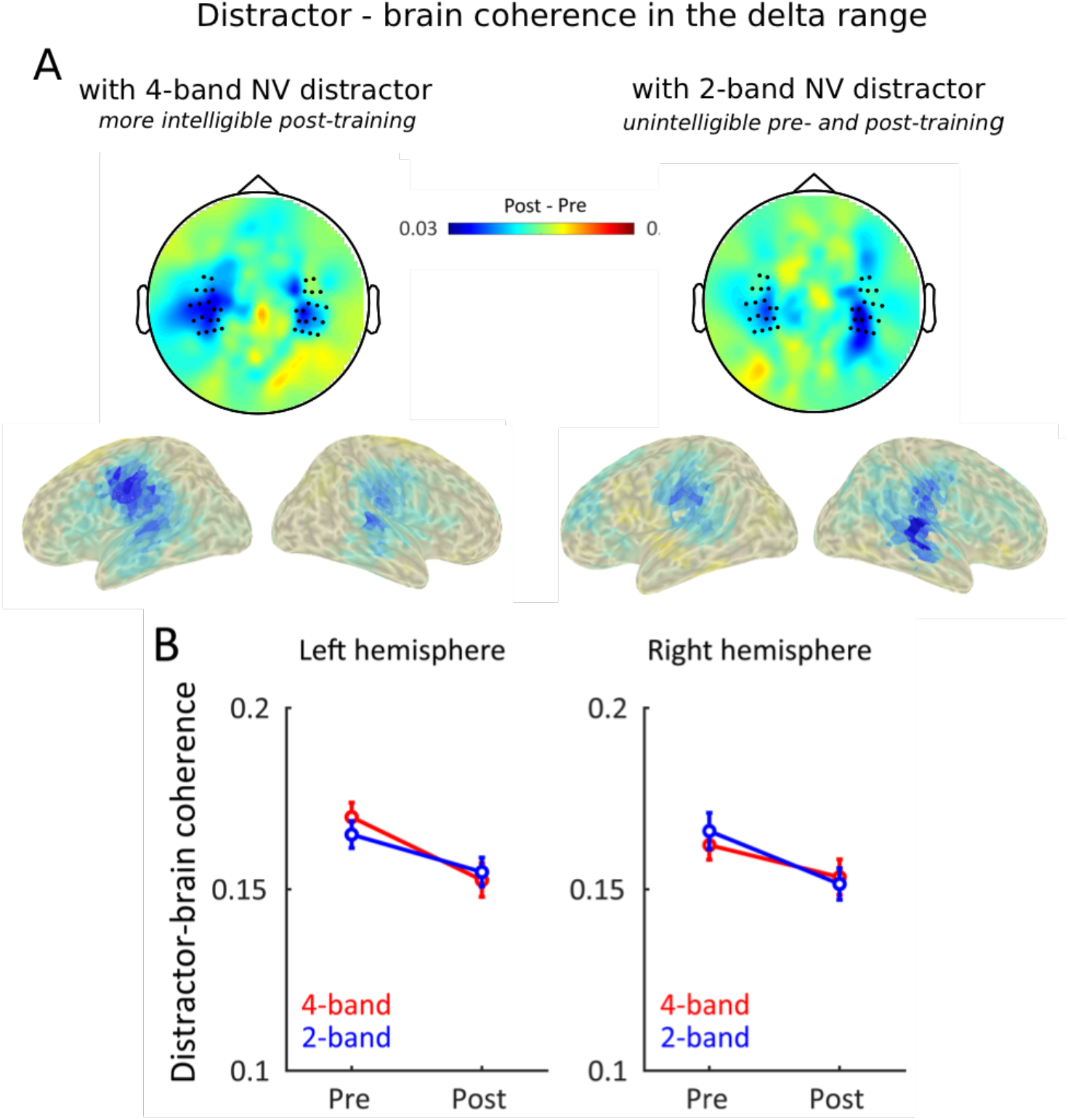
Delta tracking of distracting speech was not significantly modulated by its intelligibility. (A) Topographies and source reconstruction of changes in delta tracking (Post-training minus Pre-training). Delta tracking of distracting speech reduced after training, this irrespectively of the distracting signal. (B) Delta tracking averaged across selected channels in each hemisphere, when in competition with distracting 4-band NV speech (red) or 2-band NV speech (blue). Error bars indicated standard error of the mean.

Similarly, theta tracking of distracting speech was not modulated via training in our experiment (interaction NV* time: *F*(1, 24) = 0.01, *p* = 0.94, *η2* = 0.00), though it was strongly influenced by the acoustic properties of the signal (main effect of NV: *F*(1, 24) = 10.66, *p* = 0.003, *η2* = 0.31, Fig. 7). These results suggest that distractor speech-brain coherence is influenced by the distractor speech’s acoustic characteristics, but not by its intelligibility.

**Figure 7.**
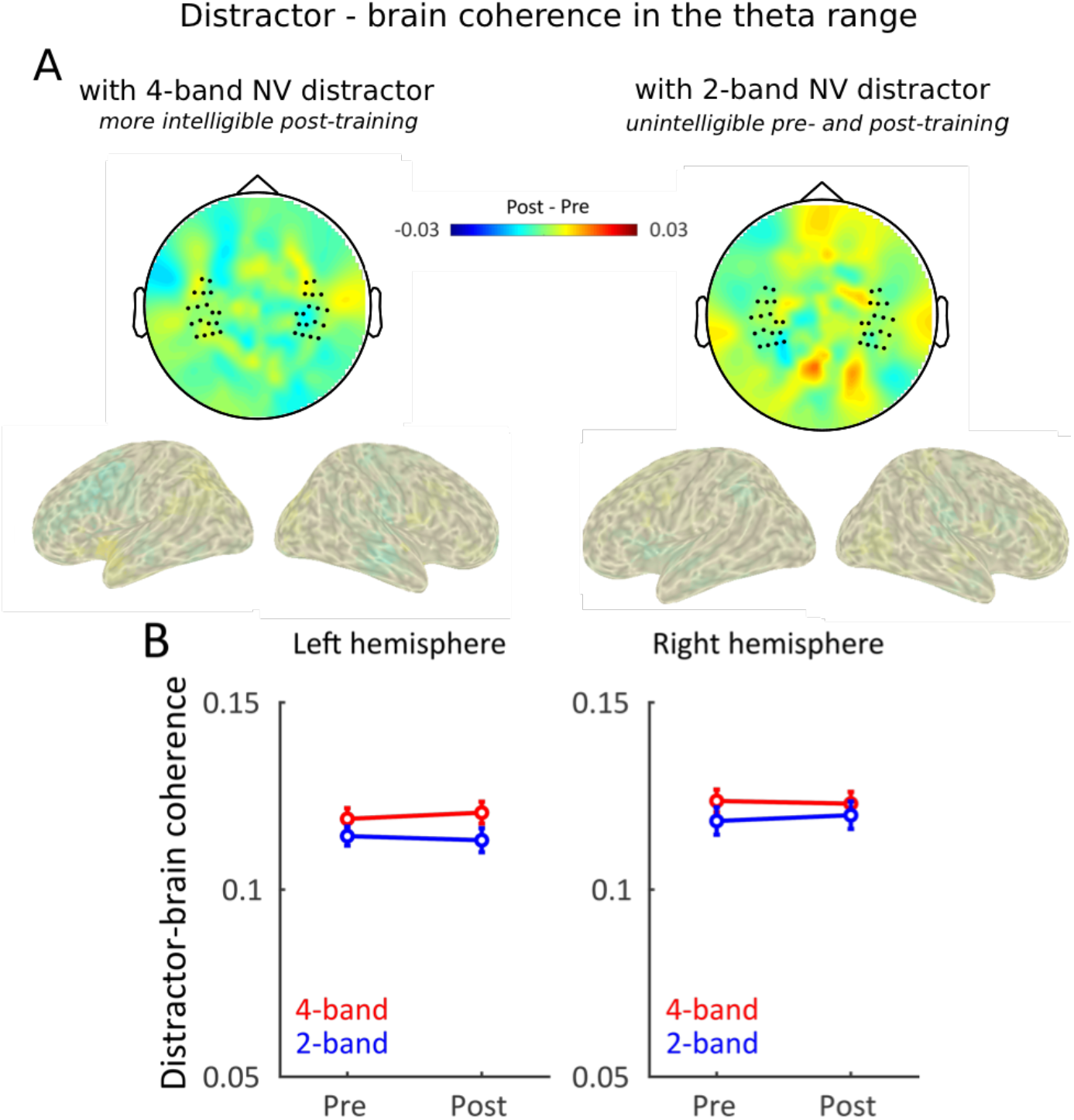
Theta tracking of distracting speech was not significantly modulated by its intelligibility. (A) Topographies and source reconstruction of changes in theta tracking (Post-training minus Pre-training). No significant changes of training were observed for both 4- and 2-band NV conditions. (B) Theta tracking averaged across selected channels in each hemisphere, when in competition with distracting 4-band NV speech (red) or 2-band NV speech (blue). Error bars indicated standard error of the mean.

## Discussion

In this MEG study, we were able to separate the linguistic and acoustic components of the masking effect between two speech signals. The main findings show that distracting speech can exert stronger interference on the neural processing of target speech when it becomes more intelligible. Crucially, we report that the presence of an intelligible speech distractor was associated with reduced neural tracking of target speech in auditory areas, even though neural tracking of the distractor was not significantly affected by its level of intelligibility. Altered tracking of target speech could represent a crucial influence on its comprehension when it is heard together with intelligible distractor speech. Moreover, neural oscillations at multiple time scales likely played different roles during speech processing: the neural tracking of target speech reduced in the presence of an intelligible distractor in the delta range (1–4 Hz) but did not do so in the theta range (4–8 Hz).

Since the classic work on the cocktail-party problem 60 years ago (Cherry, 1953), researchers have put a lot of effort into understanding the competing processing of target speech and background signals (Brungart et al., 2001; Dai et al., 2017, 2018, Ding and Simon, 2012a, 2012b; Mesgarani and Chang, 2012; Scott et al., 2009, 2004; Zion Golumbic et al., 2013). It has been suggested that distracting signals exert influence on understanding target speech depending on their intelligibility. For example, researchers have shown that speech signals impair comprehension more strongly than unintelligible sounds (Brouwer et al., 2012; Brungart et al., 2001; Garadat et al., 2009; Hygge Staffan et al., 1992; Rhebergen et al., 2005; Scott et al., 2004). However, the previous studies often manipulated the intelligibility of distracting sounds that typically affect both acoustic and linguistic content (e.g. speech *vs.* reversed speech, or native *vs.* non-native speech), leaving distinctions between acoustic interference and linguistic interference unresolved. We used a training paradigm (Dai et al., 2017) which allowed us to manipulate the intelligibility of distracting speech without changing its acoustic component, and therefore isolated the higher-order linguistic competition from lower-order effects. Our results replicate previous reports and confirm that intelligible speech is a stronger distractor than unintelligible speech (Dai et al., 2017). Given the training manipulation, stronger masking is not due to the similarity of acoustic aspects between the target and the distracting speech. Instead, it reflects effects of the higher-order linguistic information that can be extracted from the distracting signals after training. These results certainly do not exclude the possibility that acoustic information in distracting speech can have a masking effect. Rather, they suggest that acoustic masking is only part of the story, and linguistic masking offers extra interference.

Our results show that the neural tracking of target speech in the delta range reduced with a more intelligible distracting speech. In multi-speech scenes, both the target and distracting speech have multi-level information ranging from their acoustic features to linguistic meanings, and therefore their competition could happen on each level of the processing hierarchy. Based on our design, we argue that the changes we observe in target speech-brain tracking could not be attributed to acoustic components, but are rather the consequence of linguistic competition between distractor and target speech. Hence, we interpret the reduction of delta tracking of target speech as the degradation of its linguistic processing due to the competition of concurrent linguistic information. This is consistent with recent reports showing that speech-brain tracking in the delta range has been linked to the encoding of linguistic information (Di Liberto et al., 2018; Ding et al., 2016), and with neural frameworks of speech analysis suggesting that higher-level linguistic processing involves neural oscillations within the delta range (Kösem and Wassenhove, 2017; Luo and Poeppel, 2012; Poeppel, 2014).

In line with previous reports, we show that the brain concurrently tracks the slow dynamics of both attended and ignored speech (Ding and Simon, 2012a; Fiedler et al., 2019; Zion Golumbic et al., 2013). However, in apparent contradiction with the target speech-brain tracking results, the neural tracking of distracting signals was not significantly modulated by its intelligibility. The link between strength of entrainment and intelligibility is debated due to the contradictory findings: studies have reported that low-frequency speech – brain tracking is stronger when speech is intelligible, in single speaker (Ahissar et al., 2001; Ding and Simon, 2013; Doelling et al., 2014; Peelle et al., 2012; Zoefel et al., 2018) and in cocktail party settings (Ding et al., 2014; Riecke et al., 2018), while other studies failed to find a correlation between speech-brain tracking and intelligibility (Howard and Poeppel, 2010; Millman et al., 2014; Peña and Melloni, 2012; Zoefel and VanRullen, 2016). A likely source of the different results is acoustic confounds (Kösem and Wassenhove, 2017). Here, we carefully controlled for acoustic confounds and did not find significant evidence that neural tracking of distracting signals increased in strength with its intelligibility. Our results still show that distracting linguistic information is processed to some extent by the brain, as we found that distractor intelligibility decreased the neural tracking of target speech. A first tentative explanation for the present results is that speech-brain tracking would not always reflect linguistic processing in speech. Speech-brain tracking could reflect a domain-general temporal attention mechanism (Schroeder & Lakatos, 2009) that would influence speech parsing and hence comprehension (Kösem et al., 2018; 2020; van Bree et al., 2021). An alternative explanation is that speech-brain tracking would correctly mark the processing of phrasal and sentential information in speech, and that linguistic processing of unattended speech would be limited to phonological, sub-lexical and /or lexical semantic level. Recent findings have shown that the neural tracking of larger linguistic structures (e.g. words that requires the sequential grouping of two syllables) requires attention (Ding et al., 2018). In line with this, we find that delta tracking of distractor speech is not dependent on its intelligibility, suggesting that linguistic processing of the distractor stops before the sequential grouping of syllabic/word linguistic segments. The more intelligible distractor would still be able to compete with the phonological and/or lexical processing of target speech, or to capture auditory attention normally directed toward the target, which would then reduce neural tracking of the attended speech signal.

Despite presenting the target and distracting signals in different ears, we showed that both distractor and target speech signals were processed to some extent in the ipsilateral auditory cortex. The effect of distractor intelligibility on neural tracking of target speech was observed in both hemispheres and irrespective of the ear of presentation of target and distractor speech. However, we observed an effect of the ear of presentation on behavioral performance in line with previous reports (Berlin et al., 1973; Brancucci et al., 2004; Della Penna et al., 2007; Ding and Simon, 2012b; Hiscock and Kinsbourne, 2011). An intelligible NV distractor impaired more strongly target speech comprehension when the distractor was presented to the right ear. This effect could be explained as a consequence of the right-ear advantage: speech is understood faster and more accurately when presented to the right ear than to the left ear (Berlin et al., 1973; Gerber and Goldman, 1971; Studdert-Kennedy and Shankweiler, 1970), supposedly because the signal presented to the right ear has priority access to the language-dominant processing network in the left hemisphere. In our experiment, distractor signals presented in the right ear would then presumably have the processing of their linguistic information facilitated, and this could cause stronger interference on the linguistic processing of target speech. This hypothesis would suggest that the left hemisphere has a strong role in the cocktail party effect. Yet, the laterality of the reported effects cannot fully be tested considering our dichotic listening design, which was originally chosen to focus on informational masking effects and alleviate the energetic masking of the distractors. Dichotic listening also implies complete peripheral separation of the target and distractor, which is rare in natural listening. Distractor and target speech could be presented in the two ears to overcome these two limitations, and we do expect that the current findings can be replicated in diotic listening environments.

In summary, our data provide evidence that, in a multi-talker environment, the intelligibility of distracting speech degrades the processing of the target speech, as indexed by a reduction of delta tracking of target speech, with no significant change in distractor speech-brain tracking.

## Materials and Data accessibility

Stimuli set are adapted from (Versfeld et al., 2000) and are available upon request to the authors. Data and code related to this paper are available upon request from the Donders Repository, a data archive hosted by the Donders Institute for Brain, Cognition and Behaviour.

## Acknowledgments

We would like to thank Ole Jensen for providing insightful suggestions on this work; Maarten van den Heuvel and Vera van’t Hoff for help with transcribing data. This research was supported by a Spinoza award to P.H. and by a Marie Skłodowska-Curie Individual Fellowship to A.K.

## Author contributions

B.D., A.K., and P.H. conceived and designed research; B.D. performed research; B.D. and A.K. analyzed the data; all authors contributed to interpretation of the results; B.D., A.K., J.M.M, and P. H. wrote the paper.

## Supplementary materials

**Figure S1.**
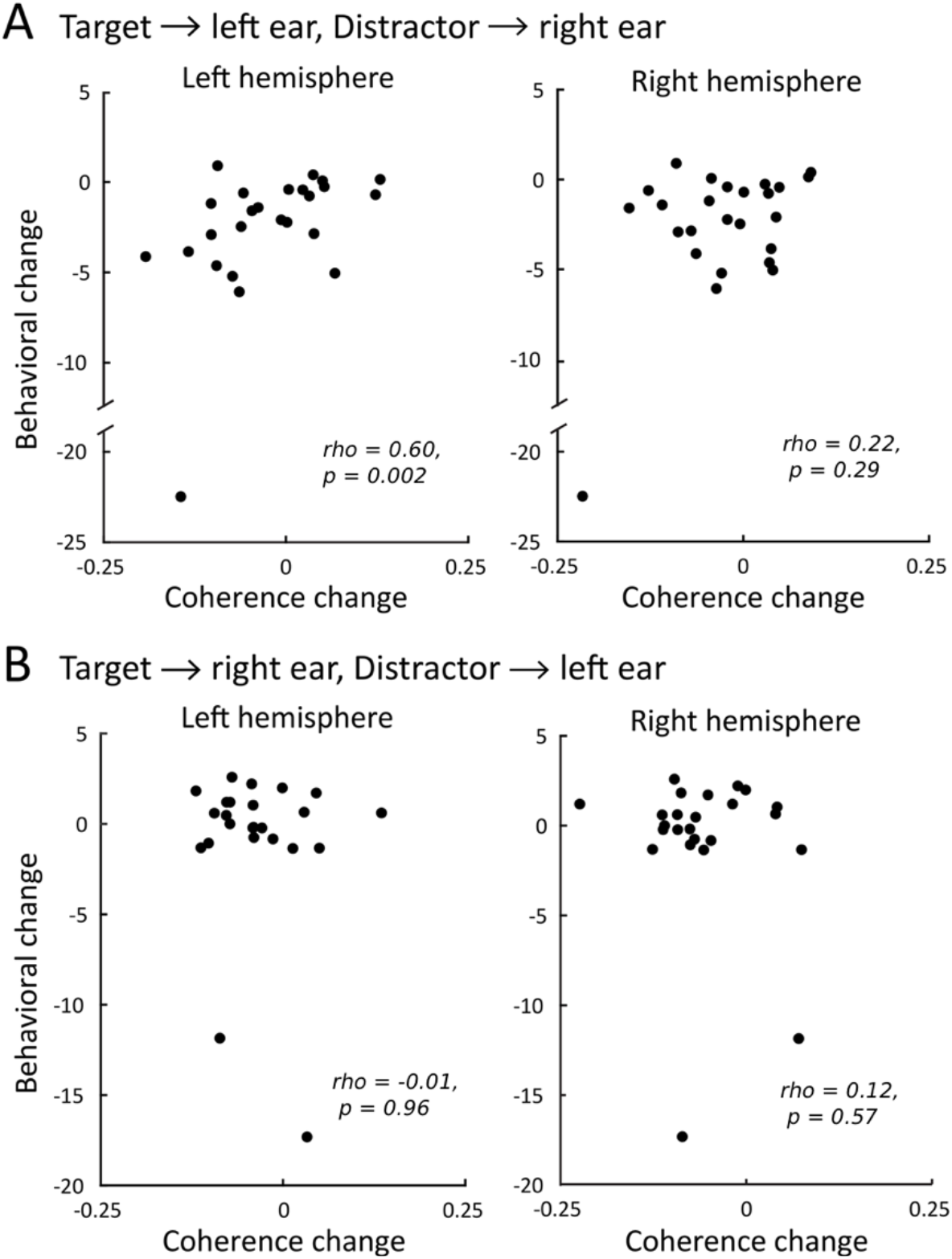
Correlation between the strength of delta tracking of target speech and the target speech intelligibility. Inter-individual correlation (Spearman’s rho) between the change in target-brain coherence in the delta range before and after training (Coherence change) and the change in target speech intelligibility before and after training, when (A) target speech was delivered into left ear; (B) Target speech was delivered into right ear. Each dot corresponds to one participant. Note that the pattern of results (significant correlation in only the top left panel) were not significant after excluding the outliers.

## References

Ahissar E, Nagarajan S, Ahissar M, Protopapas A, Mahncke H, Merzenich MM. 2001. Speech comprehension is correlated with temporal response patterns recorded from auditory cortex. Proc Natl Acad Sci 98:13367–13372. doi:10.1073/pnas.201400998

Bee MA, Micheyl C. 2008. The cocktail party problem: what is it? How can it be solved? And why should animal behaviorists study it? J Comp Psychol 122:235–51. doi:10.1037/0735-7036.122.3.235

Berlin CI, Lowe-Bell SS, Cullen JK, Thompson CL, Loovis CF. 1973. Dichotic speech perception: An interpretation of right-ear advantage and temporal offset effects. J Acoust Soc Am 53:699–709. doi:10.1121/1.1913381

Brancucci A, Babiloni C, Babiloni F, Galderisi S, Mucci A, Tecchio F, Zappasodi F, Pizzella V, Romani GL, Rossini PM. 2004. Inhibition of auditory cortical responses to ipsilateral stimuli during dichotic listening: evidence from magnetoencephalography. Eur J Neurosci 19:2329–2336. doi:10.1111/j.0953-816X.2004.03302.x

Brouwer S, Van Engen KJ, Calandruccio L, Bradlow AR. 2012. Linguistic contributions to speech-on-speech masking for native and non-native listeners: Language familiarity and semantic content. J Acoust Soc Am 131:1449–1464. doi:10.1121/1.3675943

Brungart DS. 2001. Informational and energetic masking effects in the perception of two simultaneous talkers. J Acoust Soc Am 109:1101–1109. doi:10.1121/1.1345696

Brungart DS, Simpson BD, Ericson MA, Scott KR. 2001. Informational and energetic masking effects in the perception of multiple simultaneous talkers. J Acoust Soc Am 110:2527–38.

Cherry EC. 1953. Some experiments on the recognition of speech, with one and with two ears. J Acoust Soc Am 25:975–979.

Dai B, Chen C, Long Y, Zheng L, Zhao H, Bai X, Liu W, Zhang Y, Liu L, Guo T, Ding G, Lu C. 2018. Neural mechanisms for selectively tuning in to the target speaker in a naturalistic noisy situation. Nat Commun 9:2405. doi:10.1038/s41467-018-04819-z

Dai B, McQueen JM, Hagoort P, Kösem A. 2017. Pure linguistic interference during comprehension of competing speech signals. J Acoust Soc Am 141:EL249–EL254. doi:10.1121/1.4977590

Davis MH, Johnsrude IS, Hervais-Adelman A, Taylor K, McGettigan C. 2005. Lexical information drives perceptual learning of distorted speech: evidence from the comprehension of noise-vocoded sentences. J Exp Psychol Gen 134:222–241. doi:10.1037/0096-3445.134.2.222

Della Penna S, Brancucci A, Babiloni C, Franciotti R, Pizzella V, Rossi D, Torquati K, Rossini PM, Romani GL. 2007. Lateralization of Dichotic Speech Stimuli is Based on Specific Auditory Pathway Interactions: Neuromagnetic Evidence. Cereb Cortex 17:2303–2311. doi:10.1093/cercor/bhl139

Di Liberto GM, Lalor EC, Millman RE. 2018. Causal cortical dynamics of a predictive enhancement of speech intelligibility. NeuroImage 166:247–258. doi:10.1016/j.neuroimage.2017.10.066

Di Liberto GM, O’Sullivan JA, Lalor EC. 2015. Low-Frequency Cortical Entrainment to Speech Reflects Phoneme-Level Processing. Curr Biol 25:2457–2465. doi:10.1016/j.cub.2015.08.030

Ding N, Chatterjee M, Simon JZ. 2014. Robust cortical entrainment to the speech envelope relies on the spectro-temporal fine structure. NeuroImage 88:41–46. doi:10.1016/j.neuroimage.2013.10.054

Ding N, Melloni L, Zhang H, Tian X, Poeppel D. 2016. Cortical tracking of hierarchical linguistic structures in connected speech. Nat Neurosci 19:158–164. doi:10.1038/nn.4186

Ding N, Pan X, Luo C, Su N, Zhang W, Zhang J. 2018. Attention Is Required for Knowledge-Based Sequential Grouping: Insights from the Integration of Syllables into Words. J Neurosci 38:1178–1188. doi:10.1523/JNEUROSCI.2606-17.2017

Ding N, Simon JZ. 2014. Cortical entrainment to continuous speech: functional roles and interpretations. Front Hum Neurosci 8. doi:10.3389/fnhum.2014.00311

Ding N, Simon JZ. 2013. Adaptive temporal encoding leads to a background-insensitive cortical representation of speech. J Neurosci 33:5728–35. doi:10.1523/JNEUROSCI.5297-12.2013

Ding N, Simon JZ. 2012a. Emergence of neural encoding of auditory objects while listening to competing speakers. Proc Natl Acad Sci U A 109:11854–9. doi:10.1073/pnas.1205381109

Ding N, Simon JZ. 2012b. Neural coding of continuous speech in auditory cortex during monaural and dichotic listening. J Neurophysiol 107:78–89. doi:10.1152/jn.00297.2011

Doelling KB, Arnal LH, Ghitza O, Poeppel D. 2014. Acoustic landmarks drive delta–theta oscillations to enable speech comprehension by facilitating perceptual parsing. NeuroImage 85, Part 2:761–768. doi:10.1016/j.neuroimage.2013.06.035

Ellermeier W, Kattner F, Ueda K, Doumoto K, Nakajima Y. 2015. Memory disruption by irrelevant noise-vocoded speech: Effects of native language and the number of frequency bands. J Acoust Soc Am 138:1561–1569. doi:10.1121/1.4928954

Evans S, Davis MH. 2015. Hierarchical Organization of Auditory and Motor Representations in Speech Perception: Evidence from Searchlight Similarity Analysis. Cereb Cortex 25:4772–4788. doi:10.1093/cercor/bhv136

Fiedler L, Wöstmann M, Herbst SK, Obleser J. 2019. Late cortical tracking of ignored speech facilitates neural selectivity in acoustically challenging conditions. NeuroImage 186:33–42. doi:10.1016/j.neuroimage.2018.10.057

Garadat SN, Litovsky RY, Yu G, Zeng F-G. 2009. Role of binaural hearing in speech intelligibility and spatial release from masking using vocoded speech. J Acoust Soc Am 126:2522–2535. doi:10.1121/1.3238242

Gerber SE, Goldman P. 1971. Ear Preference for Dichotically Presented Verbal Stimuli as a Function of Report Strategies. J Acoust Soc Am 49:1163–1168. doi:10.1121/1.1912478

Giraud A-L, Poeppel D. 2012. Cortical oscillations and speech processing: emerging computational principles and operations. Nat Neurosci 15:511–517. doi:10.1038/nn.3063

Gross J, Hoogenboom N, Thut G, Schyns P, Panzeri S, Belin P, Garrod S. 2013. Speech Rhythms and Multiplexed Oscillatory Sensory Coding in the Human Brain. PLoS Biol 11:e1001752. doi:10.1371/journal.pbio.1001752

Hiscock M, Kinsbourne M. 2011. Attention and the right-ear advantage: What is the connection? Brain Cogn, Dichotic Listening Anniversary Special Issue 76:263–275. doi:10.1016/j.bandc.2011.03.016

Hoen M, Meunier F, Grataloup C-L, Pellegrino F, Grimault N, Perrin F, Perrot X, Collet L. 2007. Phonetic and lexical interferences in informational masking during speech-in-speech comprehension. Speech Commun 49:905–916. doi:10.1016/j.specom.2007.05.008

Howard MF, Poeppel D. 2010. Discrimination of speech stimuli based on neuronal response phase patterns depends on acoustics but not comprehension. J Neurophysiol 104:2500–11. doi:10.1152/jn.00251.2010

Hygge Staffan, Rönnberg Jerker, Larsby Birgitta, Arlinger Stig. 1992. Normal-Hearing and Hearing-Impaired Subjects’ Ability to Just Follow Conversation in Competing Speech, Reversed Speech, and Noise Backgrounds. J Speech Lang Hear Res 35:208–215. doi:10.1044/jshr.3501.208

Kösem A, Basirat A, Azizi L, Wassenhove V van. 2016. High-frequency neural activity predicts word parsing in ambiguous speech streams. J Neurophysiol 116:2497–2512. doi:10.1152/jn.00074.2016

Kösem, A., Bosker, H. R., Takashima, A., Meyer, A., Jensen, O., & Hagoort, P. 2018. Neural entrainment determines the words we hear. Cur Biol, 28: 2867–2875.

Kösem, A., Bosker, H. R., Jensen, O., Hagoort, P., & Riecke, L. 2020. Biasing the perception of spoken words with transcranial alternating current stimulation. J Cog Neurosci, 32: 1428–1437.

Kösem A, Wassenhove V van. 2017. Distinct contributions of low- and high-frequency neural oscillations to speech comprehension. Lang Cogn Neurosci 32:536–544. doi:10.1080/23273798.2016.1238495

Lakatos P, Karmos G, Mehta AD, Ulbert I, Schroeder CE. 2008. Entrainment of Neuronal Oscillations as a Mechanism of Attentional Selection. Science 320:110–113. doi:10.1126/science.1154735

Luo H, Poeppel D. 2012. Cortical oscillations in auditory perception and speech: evidence for two temporal windows in human auditory cortex. Front Psychol 3:170. doi:10.3389/fpsyg.2012.00170

Luo H, Poeppel D. 2007. Phase Patterns of Neuronal Responses Reliably Discriminate Speech in Human Auditory Cortex. Neuron 54:1001–1010. doi:10.1016/j.neuron.2007.06.004

Mattys SL, Davis MH, Bradlow AR, Scott SK. 2012. Speech recognition in adverse conditions: A review. Lang Cogn Process 27:953–978. doi:10.1080/01690965.2012.705006

Mesgarani N, Chang EF. 2012. Selective cortical representation of attended speaker in multi-talker speech perception. Nature 485:233–6. doi:10.1038/nature11020

Millman RE, Johnson SR, Prendergast G. 2014. The Role of Phase-locking to the Temporal Envelope of Speech in Auditory Perception and Speech Intelligibility. J Cogn Neurosci 27:533–545. doi:10.1162/jocn_a_00719

Obleser, J., & Kayser, C. (2019). Neural entrainment and attentional selection in the listening brain. Trends in cognitive sciences, 23(11), 913–926.

Oever S ten, Schroeder CE, Poeppel D, Atteveldt N van, Mehta AD, Mégevand P, Groppe DM, Zion-Golumbic E. 2017. Low-Frequency Cortical Oscillations Entrain to Subthreshold Rhythmic Auditory Stimuli. J Neurosci 37:4903–4912. doi:10.1523/JNEUROSCI.3658-16.2017

Oostenveld R, Fries P, Maris E, Schoffelen J-M. 2011. FieldTrip: Open Source Software for Advanced Analysis of MEG, EEG, and Invasive Electrophysiological Data. Intell Neurosci 2011:1:1–1:9. doi:10.1155/2011/156869

Park H, Ince RAA, Schyns PG, Thut G, Gross J. 2015. Frontal Top-Down Signals Increase Coupling of Auditory Low-Frequency Oscillations to Continuous Speech in Human Listeners. Curr Biol 25:1649–1653. doi:10.1016/j.cub.2015.04.049

Peelle JE, Gross J, Davis MH. 2012. Phase-Locked Responses to Speech in Human Auditory Cortex are Enhanced During Comprehension. Cereb Cortex 23:1378–1387. doi:10.1093/cercor/bhs118

Peña M, Melloni L. 2012. Brain Oscillations during Spoken Sentence Processing. J Cogn Neurosci 24:1149–1164. doi:10.1162/jocn_a_00144

Poeppel D. 2014. The neuroanatomic and neurophysiological infrastructure for speech and language. Curr Opin Neurobiol, SI: Communication and language 28:142–149. doi:10.1016/j.conb.2014.07.005

Rhebergen KS, Versfeld NJ, Dreschler WA. 2005. Release from informational masking by time reversal of native and non-native interfering speech. J Acoust Soc Am 118:1274–1277. doi:10.1121/1.2000751

Riecke L, Formisano E, Sorger B, Başkent D, Gaudrain E. 2018. Neural Entrainment to Speech Modulates Speech Intelligibility. Curr Biol 28:161–169.e5. doi:10.1016/j.cub.2017.11.033

Schroeder, C. E., & Lakatos, P. 2009. Low-frequency neuronal oscillations as instruments of sensory selection. Trends Neurosci, 32:9–18.

Scott SK, Rosen S, Beaman CP, Davis JP, Wise RJS. 2009. The neural processing of masked speech: Evidence for different mechanisms in the left and right temporal lobes. J Acoust Soc Am 125:1737–1743. doi:10.1121/1.3050255

Scott SK, Rosen S, Wickham L, Wise RJS. 2004. A positron emission tomography study of the neural basis of informational and energetic masking effects in speech perception. J Acoust Soc Am 115:813. doi:10.1121/1.1639336

Shannon RV, Zeng FG, Kamath V, Wygonski J, Ekelid M. 1995. Speech recognition with primarily temporal cues. Science 270:303–4.

Sohoglu E, Davis MH. 2016. Perceptual learning of degraded speech by minimizing prediction error. Proc Natl Acad Sci 113:E1747–E1756. doi:10.1073/pnas.1523266113

Studdert-Kennedy M, Shankweiler D. 1970. Hemispheric Specialization for Speech Perception. J Acoust Soc Am 48:579–594. doi:10.1121/1.1912174

van Bree, S., Sohoglu, E., Davis, M. H., & Zoefel, B. 2021. Sustained neural rhythms reveal endogenous oscillations supporting speech perception. PLoS biology, 19: e3001142.

Versfeld NJ, Daalder L, Festen JM, Houtgast T. 2000. Method for the selection of sentence materials for efficient measurement of the speech reception threshold. J Acoust Soc Am 107:1671–1684. doi:10.1121/1.428451

Woods KJP, McDermott JH. 2015. Attentive Tracking of Sound Sources. Curr Biol 25:2238–2246. doi:10.1016/j.cub.2015.07.043

Wöstmann M, Obleser J. 2016. Acoustic Detail But Not Predictability of Task-Irrelevant Speech Disrupts Working Memory. Front Hum Neurosci 10. doi:10.3389/fnhum.2016.00538

Zion Golumbic EM, Ding N, Bickel S, Lakatos P, Schevon CA, McKhann GM, Goodman RR, Emerson R, Mehta AD, Simon JZ, Poeppel D, Schroeder CE. 2013. Mechanisms underlying selective neuronal tracking of attended speech at a “cocktail party.” Neuron 77:980–91. doi:10.1016/j.neuron.2012.12.037

Zoefel B, Archer-Boyd A, Davis MH. 2018. Phase Entrainment of Brain Oscillations Causally Modulates Neural Responses to Intelligible Speech. Curr Biol 28:401–408.e5. doi:10.1016/j.cub.2017.11.071

Zoefel B, VanRullen R. 2016. EEG oscillations entrain their phase to high-level features of speech sound. NeuroImage 124:16–23. doi:10.1016/j.neuroimage.2015.08.054

